# Water diviners multiplied: cryptic diversity in the *Niphargus aquilex* species complex in Northern Europe

**DOI:** 10.1101/2023.08.13.553147

**Authors:** Dieter Weber, Traian Brad, Alexander Weigand, Jean-François Flot

## Abstract

As for many other groups, patterns of biodiversity for subterranean crustaceans in Europe indicate larger morphospecies ranges at higher latitudes (the so-called Rapoport effect). However, this observed correlation may be artefactual if many of those high-latitude, widely distributed morphospecies are actually species complexes. To test this hypothesis, we looked for previously undetected species boundaries within *Niphargus aquilex* and *Niphargus schellenbergi*, two closely related morphospecies of groundwater amphipods widely distributed in northern Europe, by sequencing fragments of the mitochondrial cytochrome *c* oxidase subunit I gene (COI) and of the nuclear 28S ribosomal RNA gene of 198 individuals collected across their area of distribution. Distance-based and allele sharing-based species delimitation approaches were congruent in revealing the existence of at least eight species within *N. aquilex* and at least two species with *N. schellenbergi*. Our data demonstrate that these two common morphospecies with large ranges are actually complexes of species with narrower distributions, suggesting that the Rapoport effect might be the result of increased morphological stasis at high latitudes rather than actual differences in sizes of distribution ranges.

## Introduction

Many groups of organisms have been reported to display latitudinal gradients in ecological ranges, which higher latitudes having more species more broadly distributed than locations closer to the equator - the so called ‘Rapoport’s rule’ (1) or Rapoport effect. However, such pattern could be the result of a methodological artefact (2) or of incorrect taxonomy, for instance if high-latitude morphospecies contain more often undetected species boundaries than low-latitude morphospecies (3). A good model group to test this hypothesis is subterranean crustaceans in Europe, such as *Niphargus* amphipods. *Niphargus* is by far the most species-rich genus of freshwater amphipods, comprising hundreds of species (4) spread over Europe (except its northern part) as well as part of Central Asia and the Middle East, from Ireland to the West (5) till Kazakhstan (6) and Iran (7, 8) to the East. Niphargid amphipods exhibit a typical stygomorphic habitus, being depigmented and eyeless (9).

The taxonomy of *Niphargus* has been in flux ever since this genus was erected in 1849 (10, 11), and one of the most taxonomically troublesome part of the genus has been the so-called *Niphargus aquilex* species complex (12), which comprises currently two morphospecies: *Niphargus aquilex* (“aquilex” meaning “water diviner” in Latin) and *Niphargus schellenbergi*, both reported in the literature from rather large geographical areas (Figures 1 and 2). Although these two species are closely related, a previous study concluded that they can be told apart morphologically as well as using DNA sequences (13).

**Fig. 1.**
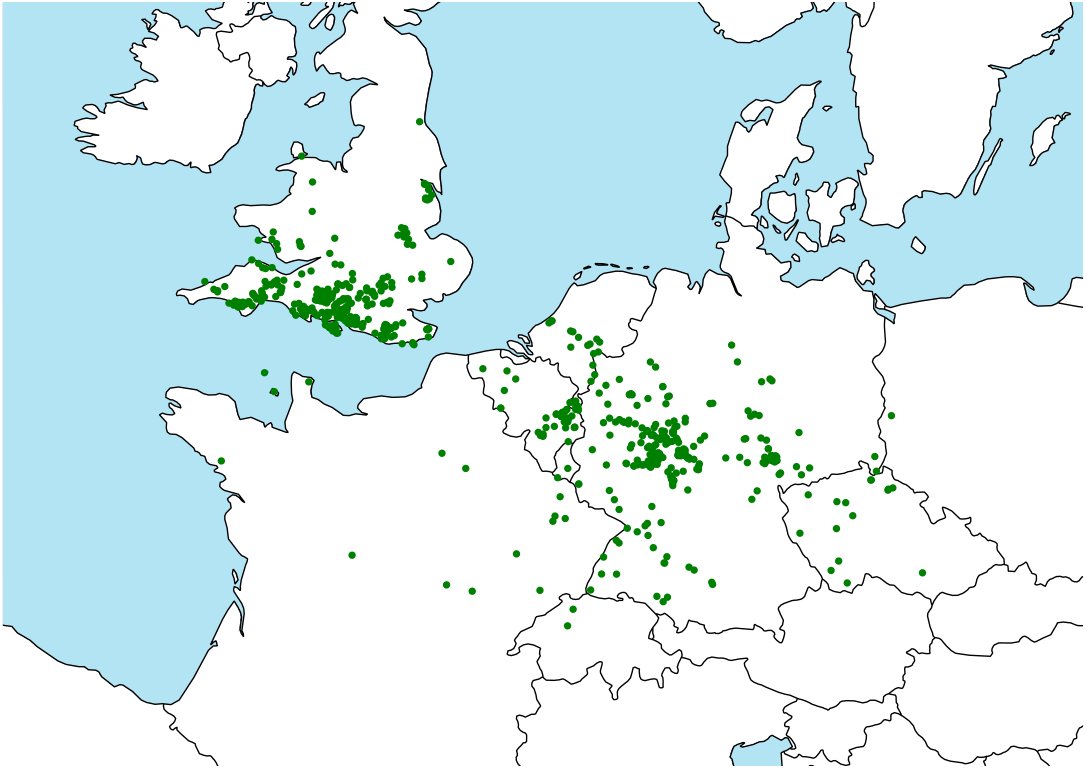
*Niphargus aquilex* (753 data points) based on published data (mainly morphological determination), including unpublished morphological data of S. Zaenker. It may include doubtful determination results.

**Fig. 2.**
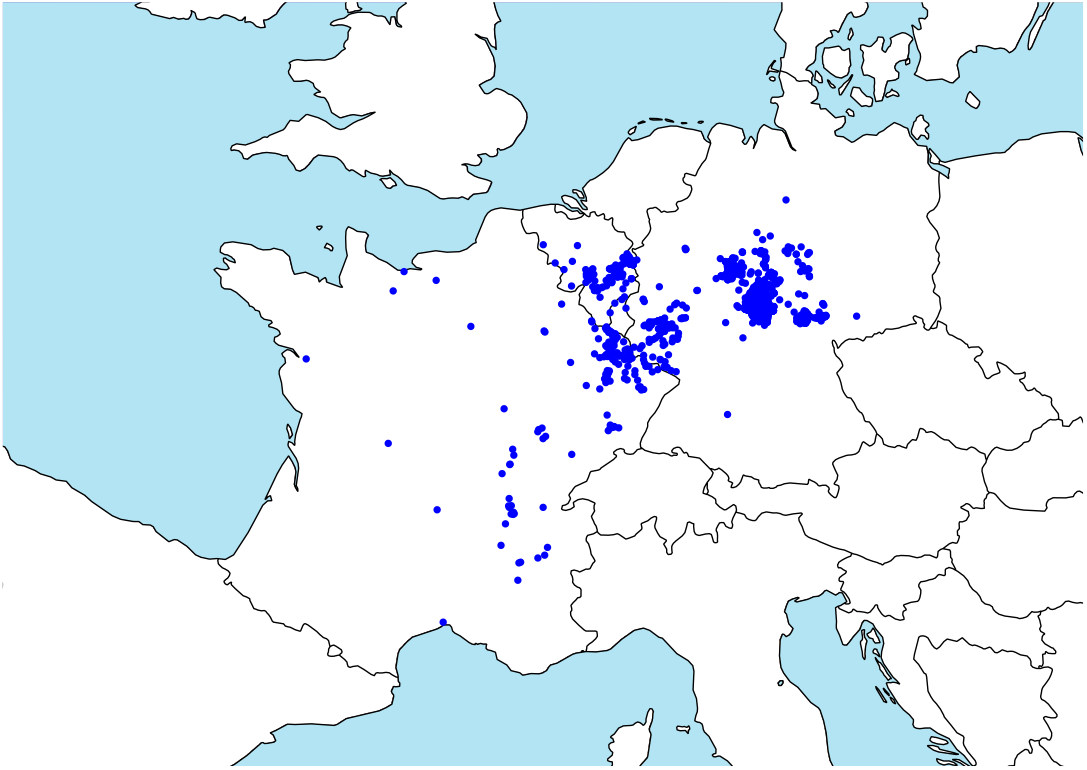
*Niphargus schellenbergi* (1892 data points) based on published data (mainly morphological determination), including unpublished morphological data of S. Zaenker. It may include doubtful determination results.

*N. aquilex* was originally described from England. The genus *Niphargus* from England was firstly mentioned as *Niphargus stygius* in 1953 by Westwood (14). Two years later, Schiödte corrected Westwood’s determination and described *N. aquilex* from Maidenhead close to London (15). 125 years later, *N. aquilex* was re-described on new material from Moretonhampstead (UK, Devon, WGS 84: Lat.: 50.7, Lon.: -3.8), and Crowborough (UK, East Sussex, Lat.: 51.1, Lon.: 0.2), with Crowborough designated as neotype locality (16); although Crowborough is a city in South East England with an area of 14 km^2^, a more precise location was not specified. *Niphargus schellenbergi* was first described in 1932 based on 5 localities with the Gännsbrünnle south of Sendelbach, Bavaria (Germany) as type locality (17). In the same year, *N. schellenbergi* was transferred to *N. aquilex* as a subspecies (18). The following year, *N. aquilex schellenbergi* was synonymised with *N. puteanus* by Schellenberg (19, 20) but subsequently the same author separated them again (21, 22). Only in 1972, 40 years after its first description, was *N. schellenbergi* erected again as a separate species distinct from *N. aquilex* (23).

The geographical spread of these two high-latitude morphospecies (Figures 1 and 2) is much larger than for *Niphargus* species from southern locations in Europe, which are generally much more localized (e.g. (24)). We therefore embarked on a molecular study aimed at testing the hypothesis that they are actually complexes of cryptic (i.e., morphologically identical) or pseudocryptic (i.e., morphologicall very similar) species with smaller ranges, as previously reported for other widespread *Niphargus* morphospecies (3, 25).

## Materials and Methods

*Niphargus* specimens for the present study were searched for between 2014 and 2018 at 17 sites in the United Kingdom, 223 in France, 5 in the Netherlands, 83 in Belgium, 121 in Luxembourg, 497 in Germany, 2 in Poland and 10 in Czechia. We collected niphargids in natural caves, abandoned mines, abandoned railway tunnels, other artificial caverns, springs, wells and the interstitial. For springs and some caves we used a set of four sieves (5 mm; 1 mm; 0.6 mm; 0.2 mm). For interstitial sites we used either the Karaman-Chappuis-method (26) or a Bou-Rouch-pump (27). In caves and artificial caverns, we collected opportunistically or using tin cans with chicken liver. In wells we used a landing net with a mesh size of 0.2 mm or occasionally tin cans with baits.

Within the research area, niphargid specimens that matched the description of the *Niphargus aquilex* species complex by Ginet (28) were selected for sequencing: coxal plate IV as high as coxal plate V and without postero-distal lobe; palma of gnathopods rectangular (not ovoid) and with 1-5 spines on the outer margin of the dactylus; uropod III of males elongated; 3rd epimeral plate with rounded postero-ventral corner or with postero-ventral corner at an angle of 90° to 100°, but not pointed; telson free of dorsal spines and as long as wide or not more than 1.2 times as long as wide.

Captured specimens were immediately preserved in 96% ethanol and kept at -20°C. Whenever possible, at least one male and one female from every site were preserved in 70% ethanol at room temperature for morphological investigation. For specimens larger than 4 mm one pereiopod was used for DNA isolation. For specimens ranging from 3 mm to 4 mm in length, two pereiopods and for specimens smaller than 3 mm the whole specimen was used. DNA was extracted following standard protocols either from the Qiagen® DNeasy Blood & Tissue Kit (Qiagen, Germany) or the NucleoSpin® Tissue Kit (Macherey-Nagel). At least one specimen per site was sequenced for two standard markers: the mitochondrial cytochrome *c* oxidase subunit I gene (COI) and the nuclear 28S ribosomal RNA gene (28S) using the same primers and PCR conditions as in (29).

Chromatograms that displayed double peaks were deconvoluted to yield phased sequences (30): briefly, the haplotypes of individuals with a single double peak were inferred directly (trivial case), those of length-variant individuals (recognizable by their numerous double peaks) were inferred using Champuru (31, 32), and finally those of individuals with a few double peaks were inferred using SeqPHASE (33) (taking advantage of the known phases of the other individuals in the dataset). Species delimitation from DNA sequences was performed using two complementary methods: ASAP (34) (assemble species by automatic partitioning), a distancebased approach, and haplowebs (35), an allele sharing-based approach. The third main type of species delimitation methods, tree-based approaches (36), was not used here as simulation studies have revealed their tendency to oversplit datasets, yielding artefactual species boundaries (37–40). Haplotype networks (haplonets) for 28S and COI were built using the program HaplowebMaker (41), which was also used to turn the 28S haplotype network into a haplotype web (haploweb) by adding curves connecting haplotypes found co-occurring in heterozygous individuals.

All sequenced specimens from Luxembourg as well as DNA isolates obtained in the laboratories of the National Museum of Natural History Luxembourg (MNHNL) are stored in the MNHNL collections. All other DNA isolates are stored at the research unit of Evolutionary Biology and Ecology of the Université libre de Bruxelles (ULB).

## Results

198 individuals were successfully sequenced for both markers. For COI, almost no double peaks were detected except in one individual that had one clear double peak, which could be due to heteroplasmy or to the presence of a nuclear COI pseudogene amplified and sequenced at the same time as the actual COI locus (42). Both haplotypes of this individual were included in downstream analyses. Similarly, double peaks in COI chromatograms were observed previously in another amphipod genus closely related to *Niphargus* (29). For 28S, 23 out of the 198 individuals sequenced displayed double peaks indicative of heterozygosity.

ASAP analysis of COI sequences (Figure 4) yielded a most-favored partition (COI-ASAP-1, with an ASAP score of 2.5) made up of 11 putative species (shown as continuous ellipses), whereas the second most-favored partition (COI-ASAP-2, with an ASAP score of 3.5) suggested 14 putative species (shown as a combination of continuous and dotted ellipses on Figure 4). By contrast, ASAP analysis of 28S sequences (Figure 3) revealed 10 putative species (28S-ASAP, with an ASAP score of 3.5), whereas delineating haplotype-sharing groups of individuals (i.e., fields for recombination following Doyle’s terminology (43)) on the 28S haploweb (28S-haploweb) yielded 13 groups (represented by letters A, B, D, F, G, H, I, J, K, L, N, R and S on Figure 3 and listed in Table 1).

**Fig. 3.**
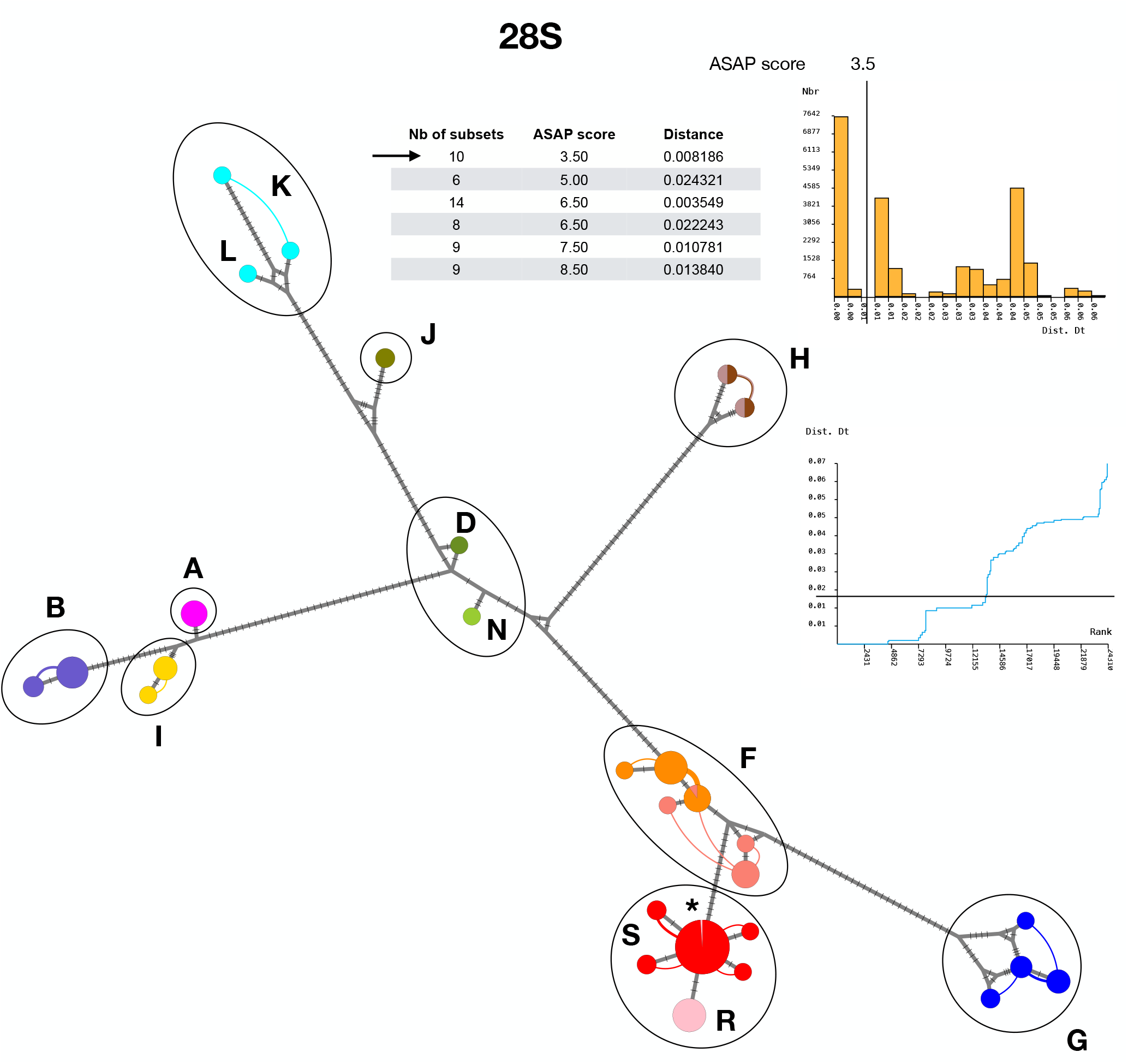
28S haploweb of the samples gathered in the present study. Curves connect haplotypes found co-occurring in heterozygous individuals.

**Fig. 4.**
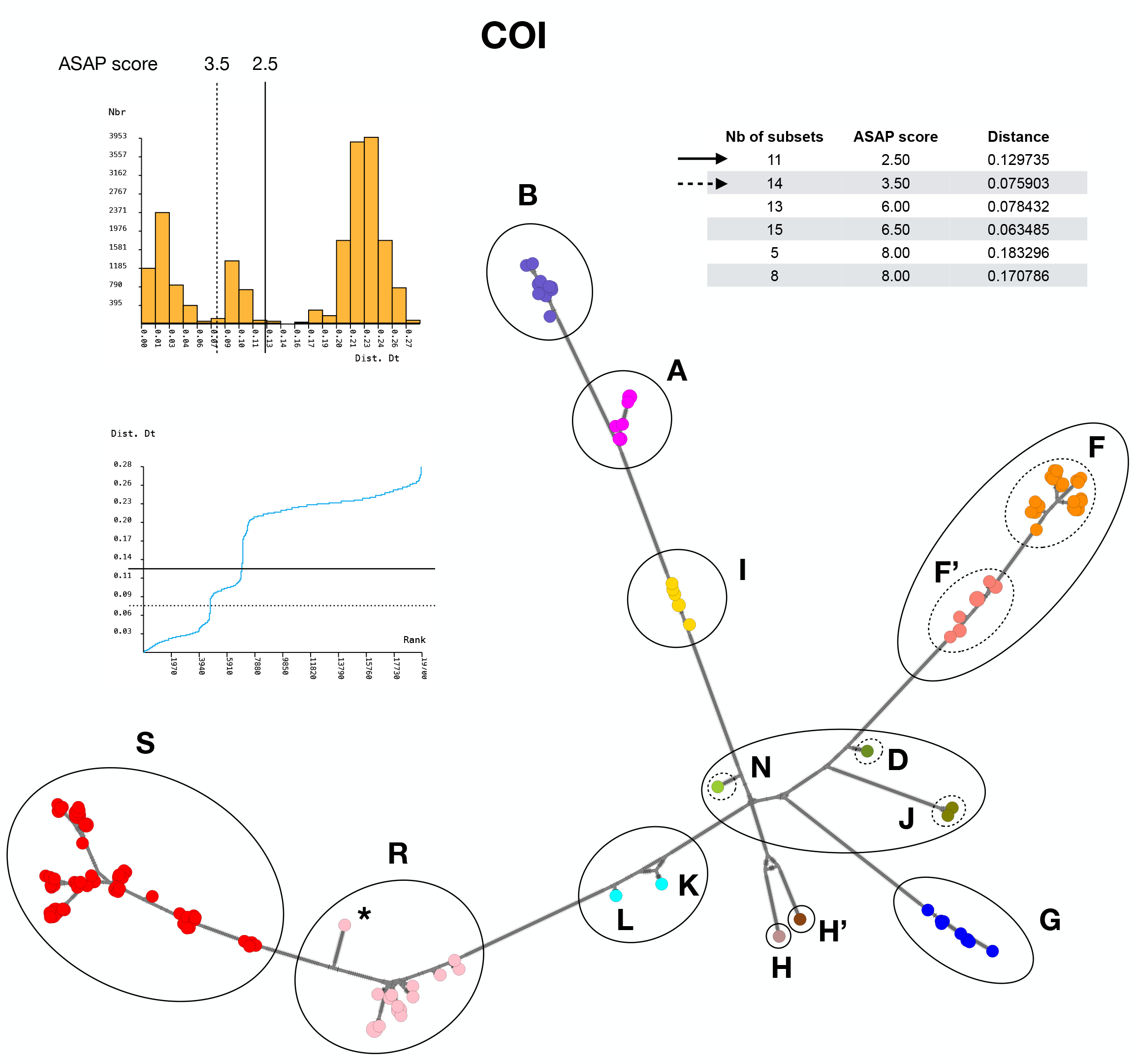
COI haplotype network of the samples gathered in the present study. Curves connect haplotypes found co-occurring in heterozygous individuals.

**Table 1.** Details of the 13 groupings inferred from 28S-haploweb.

| Group | Nb of specimens | Nb of 28S haplotypes |
| --- | --- | --- |
| A | 9 | 1 |
| B | 19 | 2 |
| D | 1 | 1 |
| F | 37 | 7 |
| G | 9 | 4 |
| H | 2 | 2 |
| I | 6 | 2 |
| J | 2 | 1 |
| K | 1 | 2 |
| L | 1 | 1 |
| N | 1 | 1 |
| R | 21 | 1 |
| S | 87 | 5 |

Among these 13 28S-haploweb groupings, four were recovered congruently by all four methods (28S-haploweb, 28S-ASAP, COI-ASAP-1 and COI-ASAP-2): these are groups A, B, H and I. Four groupings were recovered using three methods out of four: group J was supported by 28S-haploweb, 28S-ASAP and COI-ASAP-2, whereas COI-ASAP-1 lumped it with D and N; group F was supported by 28S-haplowebs, 28S-ASAP and COI-ASAP-1, but not by COI-ASAP-2 that splitted it into two (noted F and F’ on Figure 4); groups S and R were supported by 28S-haploweb, COI-ASAP-1 and COI-ASAP-2 (with the exception of one discordant individual from Chaligny (Meurthe-et-Moselle, France) that grouped with S in 28S but with R in COI; shown with an asterisk on Figures 3 and 4) but not by 28S-ASAP that lumped them together. Three groupings were recovered using two methods out of four: the distinctiveness of D and N was supported by 28S-haploweb and COI-ASAP-2, but not by 28S-ASAP (the lumped them together nor COI-ASAP-1 (that lumped them together with N); H was recovered as one putative species using both 28S-haploweb and 28S-ASAP but as two species H and H’ by COI-ASAP-1 and COI-ASAP-2. Finally, K and L were considered as two distinct species only by 28S-haploweb, whereas 28S-ASAP, COI-ASAP-1 and COI-ASAP-2 all lumped them into a single species.

## Discussion

As the congruence between the four different approaches was not perfect, the number of species in our dataset appears to lie somewhere between ten (A, B, NDJ, F, G, H, I, KL, R and S) and 15 (A, B, N, D, J, F, F’, G, H, H’, I, K, L, R and S). The intersection between these two lists comprise six groups that emerge from our analysis as strongly supported.

Groups A, B, G, I are supported by all methods and each of them correspond morphologically to *Niphargus aquilex*, confirming the existence of yet unrecognized species boundaries within this morphospecies. Group B was the only group collected at the type locality of *Niphargus aquilex*.

Groups S and R are supported by all methods except 28S distances. S corresponds to *Niphargus schellenbergi* as it includes near-topotypes of this species. As there are only two 28S mutations separating R from S (which is much less than the level of divergence within e.g. putative species G), it is not surprising that ASAP (a distance-based approach) failed to distinguish them using this marker. If a species comprises two haplotype groups that are present with the same frequency in the population, a given individual has a probability of 0.5 of having both of its haplotypes from the same group and a probability of 0.5 of having one haplotype in each group. If five individuals are randomly selected for sequencing, the probability that none of them harbor two haplotypes of different groups is 0.5^5^ hence 0.03125, which is already quite small. With 10 individuals sequenced without any “mixed-group” individual being discovered, this p-value becomes 0.5^10^ hence approximately 0.001. Of course, such calculation should take into account the respective frequencies of the two haplotype groups. In the present case, out of 108 individuals sequenced 21 individuals had both haplotypes in R and 87 had both haplotypes in S: assuming that our sampling represents approximately the respective proportions of both haplotype groups in the field, it gives a frequency of ca. 0.2 for R and ca. 0.8 for S, and the probability that we would obtain no heterozygous individual harboring one R haplotype and one S haplotype is (0.2^2^+0.8^2^)^108^=0.68^108^=8.10^−19^, a vanishingly small probability that allows us to reject the hypothesis that R and S are conspecific with extreme confidence. Still, it is striking that one individual with incongruent 28S and COI groupings was identified in a location where the distributions of the two putative species overlap: this individual could be an instance of mitochondrial DNA capture, a major source of mito-nuclear incongruence detected in many species such as chipmunks (44). Future whole-genome sequencing of the genome of this individual and comparison with those of typical R and S individuals will be required to elucidate this situation. Morphological comparison of R and S is pending, as well as the description of R as a new species distinct from (but closely related to) *Niphargus schellenbergi*.

Support for the other seven 28S-haploweb groupings is less strong and ascertaining their putative species status will require further study using more markers and/or further individual collections. Groups H and F were each split into two by COI (by both COI-ASAP-1 and COI-ASAP-2 for H, vs. only by COI-ASAP-2 for F) but not by 28S, as their 28S haplotypes were all connected and not very distant from one another. Distinct niphargid species sharing identical 28S haplotypes but distinguishable using COI as well as ITS were previously reported (45): hence, deciding whether H and F are actually single species or pairs of closely related ones will require taking a close look at additional markers. By contrast, the problem for D, J and N is the very low number of specimens sequenced so far (one for D, two for J and one for N). The situation is very similar for L and K, with only one individual sequenced for each so far for each of them. Hence, deciding whether D, J and N on the one hand and K and L on the other hand are distinct species or not will require sequencing additional samples rather than additional markers.

## Conclusions

Our study focused on two widespread *Niphargus* morphospecies emblematic of western European groundwater: *Niphargus aquilex* and *Niphargus schellenbergi*. It confirmed the distinctiveness of these two taxonomic entities but revealed cryptic species boundaries within each of the two: *Niphargus aquilex* comprises at least eight species (A, B, NDJ, F, G, H, KL, I), whereas *Niphargus schellengergi* can be split into two. This demonstrate that these two common morphospecies with large ranges are actually complexes of species with narrower distributions, suggesting that the Rapoport effect might be the result of increased morphological stasis at high latitudes rather than larger distribution ranges.

The methodology used here based on the congruence of two markers (28S and COI) analysed in two different ways (ASAP and haplowebs) appears generally effective at delimiting *Niphargus* species but future studies will benefit from replacing 28S with a longer ribosomal DNA marker: either the internal transcribed spacers (ITS) region, a marker that has shown excellent potential to delineate niphargid species in previous studies (45–53), or a larger region comprising both 28S, ITS and possibly the neighboring 18S ribosomal RNA gene to maximize phylogenetic resolution. Given that ITS in niphargid amphipods is very long and may represent the longest ITS of the whole animal kingdom (54), long-read amplicon sequencing using e.g. Nanopore or PacBio will be required if a larger region than just ITS is targeted. However, even replacing 28S with ITS (or 28S+ITS) in future studies will not solve the main challenge in *Niphargus* taxonomy: the rarity of some species, for which only one or two individuals are collected. In such cases, the usual approach is to sequence and publish data as they come but to wait for taxonomic decision until an ulterior date when more individuals may become available for study. However, this means that rare species, the one that generally require urgent protection, remain unnamed and therefore below the radars of scientists and stakeholders for long periods of time. To alleviate this problem, a better solution might be to name them and describe them provisionally, and to revise their description where more samples before available.

## ACKNOWLEDGEMENTS

JFF’s laboratory is supported by the Belgian Fond de la Recherche Scientifique - FNRS via “Projet de Recherches” grant T.00078.23. Molecular analyses at the MNHNL were financially supported by the Ministry of the Environment, Climate and Sustainable Development Luxembourg.

